# Nanobody-mediated control of gene expression and epigenetic memory

**DOI:** 10.1101/2020.09.09.290015

**Authors:** Mike V. Van, Taihei Fujimori, Lacramioara Bintu

## Abstract

Targeting chromatin regulators to specific genomic locations for gene control is emerging as a powerful method in basic research and synthetic biology. However, many chromatin regulators are large, making them difficult to deliver and combine in mammalian cells. Here, we developed a new strategy for gene control using small nanobodies that bind and recruit endogenous chromatin regulators to a gene. We show that an antiGFP nanobody can be used to simultaneously visualize GFP-tagged chromatin regulators and control gene expression, and that nanobodies against HP1 and DNMT1 can silence a reporter gene. Moreover, combining nanobodies together or with other regulators, such as DNMT3A or KRAB, can enhance silencing speed and epigenetic memory. Finally, we use the slow silencing speed and high memory of antiDNMT1 to build a signal duration timer and recorder. These results set the basis for using nanobodies against chromatin regulators for controlling gene expression and epigenetic memory.

## Introduction

Controlling gene expression levels and epigenetic memory is critical for dissecting biological processes, such as development and cancer, and for developing synthetic biology applications that rely on engineering human cells. As a result, the development of tools that can target a specific gene and alter its expression in a controlled manner is crucial for the advancement of this field. Typically, these tools consist of one or more effector domains from transcription factors or chromatin regulators, fused to a DNA binding domain that can bind to a specific sequence (e.g. TetR or Gal4, mostly used as components of synthetic genetic circuits) or be engineered to target any sequence in the genome (e.g. Zinc Fingers, TALEs, or dCas9).

In mammalian cells, the most efficient way to control gene expression and impart epigenetic memory is to engage multiple chromatin regulators (CRs) in a combinatorial fashion. For example, the Krüppel-associated box (KRAB), a strong repressive domain commonly used in gene circuits and CRISPRi applications^1^, recruits KAP1 which in turn interacts with other repressors such as histone deacetylases via the NuRD complex^2^, histone methylase SETDB1, and heterochromatic protein HP1^3^. Previous works have also shown that combining other CRs, such as MeCP2 and DNMT3A, with KRAB can further improve its silencing strength and epigenetic memory. Adding MeCP2 increases silencing strength at certain endogenous genes^4^ while adding DNMT3A can increase silencing duration after a short initial pulse of gene targeting^5–7^. Further transcriptional control via histone deacetylation^8^, acetylation^9^, methylation^10^ and demethylation^11^ by CRs fused with dCas9 has also been reported, highlighting the flexibility of these tools in imparting diverse epigenetic modifications.

Most chromatin regulators are large, consisting of multiple domains that are necessary for their function, either to stimulate catalytic function or to mediate interactions with other members of the complexes they are part of. For example, HP1’s role in the formation of transcriptionally inactive heterochromatin is dependent on its two protein domains, its chromodomain (CD) and an evolutionary related chromoshadow domain (CSD). The CD allows HP1 to bind to H3K9 methylated chromatin and is required for its gene silencing function^12,13^, while the CSD helps HP1 bind to other HP1-like proteins^14^ as well as other interactors such as SUV39H1 and DNA methyltransferases^15^ that may further enhance these silencing effects. Furthermore, sometimes the catalytic domain of a CR is not sufficient for silencing. Unlike DNMT3A and DNMT3B, the C-terminal catalytic domain of DNMT1 has been shown to be catalytically incapable of methylating DNA by itself^16,17^, requiring a large part of the N-terminal domain for its activity^18^. Therefore, in order to obtain efficient gene control, it is often desirable to recruit full-length CRs to a target gene.

Due to the large size of CRs and the requirement to use them in combinations, it is difficult to build compact gene regulatory tools that can fit in viral delivery vectors such as AAV (payload limit ≤ 4.7 kb) or lentivirus (payload limit ≤ 9.7 kb). This problem is further exacerbated by the size of *Streptococcus pyogenes* Cas9 (SpCas9), which at over 4.2 kb makes adding one or more CRs challenging. To overcome this size limit, a smaller variant of dCas9 has been engineered by deleting various functional domains; and when combined with a small transactivation domain was able to barely fit within the packaging limit of AAV and showed efficient activation activity^19^. Additionally, splitting the dCas9 protein (e.g. by utilizing two dimerizable fragments^20^ or the intein-mediated trans-splicing system^21,22^) or simply separating the dCas9 protein and guide RNAs into two different viral delivery vectors have been used to overcome this packaging limit. However, these latter approaches can suffer from poor editing efficiency due to the inability to deliver all payload components into one given cell. Therefore, while there are current efforts to find and test smaller CRISPR variants (reviewed in ^23^), it is equally important to develop smaller effector domains.

Here, we take a new approach to building smaller tools for gene regulation: instead of fusing large CRs to the DNA binding domain, we use single-domain antibodies (also called nanobodies) to recruit endogenous CRs from the existing cellular chromatin network. Due to their small size (~15 kDa), strong binding affinity, and high stability, nanobodies have been utilized successfully in many research applications, including live cell imaging, protein immunoprecipitation, and fluorescent protein enhancement (reviewed in ^24,25^). We show that nanobodies can be used to silence gene expression and impart epigenetic memory, and can enhance the function of commonly used transcriptional effectors. Using nanobodies to recruit endogenous CRs to a gene of interest for transcriptional control could offer a better opportunity to recruit entire chromatin complexes, taking advantage of all domains essential for activity and their interactions.

## Results

### Nanobodies against GFP-tagged chromatin regulators are used to control gene expression

In order to test if nanobodies can be used to recruit chromatin regulators (CRs) for efficient gene expression control, we started with a well-characterized nanobody against GFP^26–28^. We fused this nanobody to a reverse Tetracycline Repressor (rTetR) DNA-binding domain and used it to recruit various publicly available GFP-tagged CRs to a TagRFP fluorescent reporter gene located at the AAVS1 locus in HEK293T cells (Fig. 1a). This reporter contains five Tet operator (TetO) binding sites upstream of a constitutive pEF promoter driving the expression of the fluorescent gene. Upon the addition of doxycycline (dox) to the cell media, rTetR binds to these sites, allowing us to control recruitment and release of the rTetR-nanobody fusion at the reporter.

**Figure 1:**
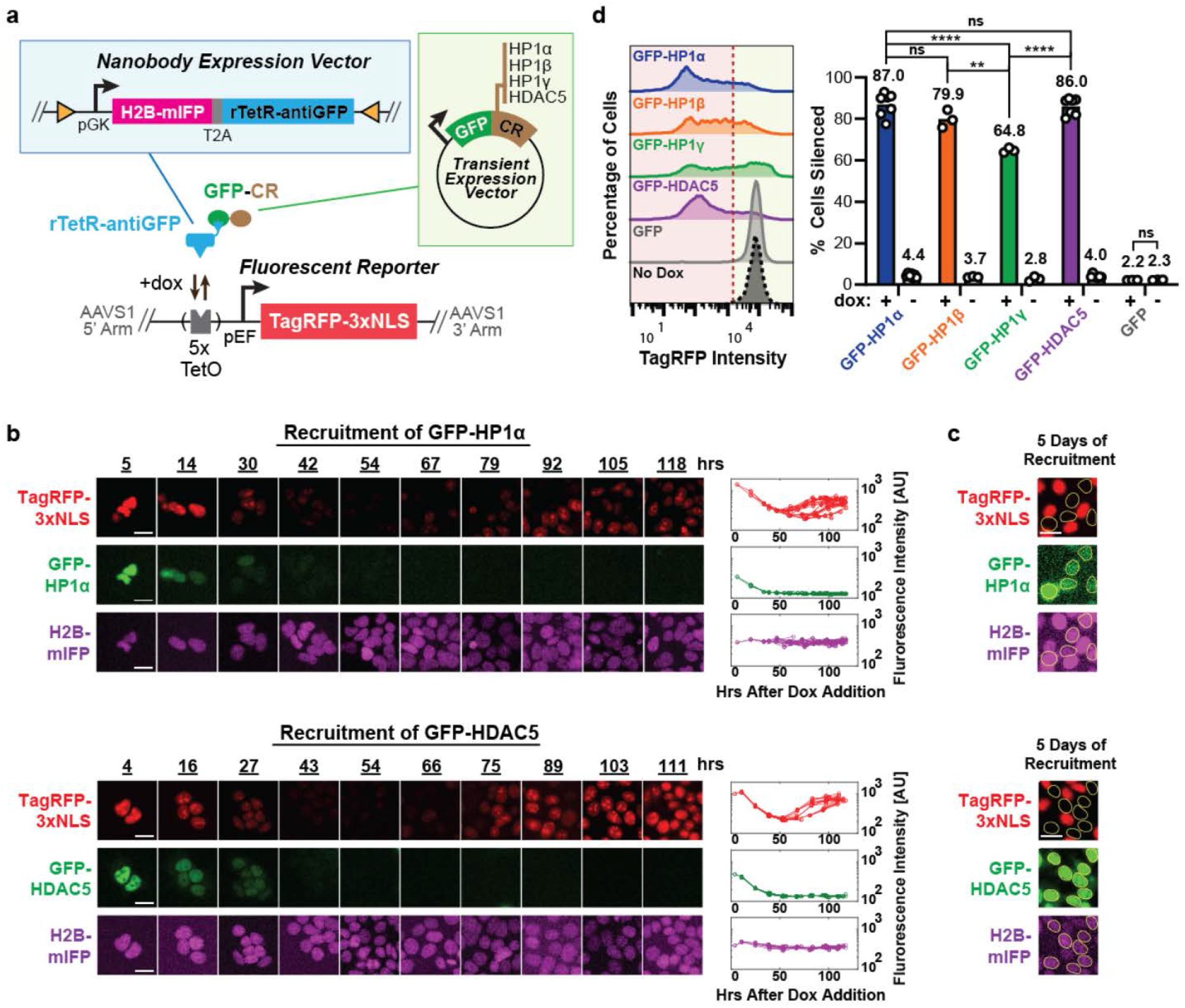
Nanobodies against GFP-tagged chromatin regulators allow for gene expression control. (**a**) Construct for constitutive coexpression (under the pGK promoter) of H2B-mIFP and antiGFP nanobody fused with the rTetR DNA binding protein (separated by the self-cleaving peptide T2A) was randomly integrated into HEK293T cells by PiggyBac (blue box, top). These cells also contain a TagRFP reporter gene integrated at the AAVS1 “safe harbor” locus and driven by the pEF promoter (bottom). Five TetO binding sites allow binding of rTetR upstream of the reporter gene upon dox addition. The nuclear localization signal (NLS) and H2B domains localize fluorescent protein signals to the nucleus, improving quantitation during time-lapse imaging. Plasmids expressing GFP-tagged CRs (HP1α, HP1β, HP1γ, and HDAC5) were transiently transfected into cells (green box, right). (**b**) Time-lapse imaging of cells upon recruitment of GFP-tagged HP1α (top) and HDAC5 (bottom). Cells stably expressing the reporter (TagRFP, red) and the rTetR-antiGFP fusion (mIFP, purple) were transiently transfected with GFP-HP1α or GFP-HDAC5 (GFP, green). Cells were treated with dox throughout the movie starting at time 0 hours. Example images across the time-lapse movie (left) and single-cell traces for individual cell lineages as they undergo up to 8 cell divisions (right). (**c**) Cells that still have GFP-CR expression (yellow circles) by day 5 of recruitment remain silenced. Bars, 20 μm. (**d**) (left) Fluorescence distributions measured by flow cytometry showing reporter silencing after recruitment of GFP-CRs (+dox) for 5 days. Cells were gated for the presence of both GFP-CR (GFP positive) and rTetR-antiGFP (mIFP positive). The red dotted line was used to determine the percentage of cells silenced shown on the right. (right) Mean percentage of cells silenced upon presence or absence of dox for 5 days. Each dot is an independently transfected biological replicate (GFP-HP1α: n = 7; GFP-HP1β: n = 3; GFP-HP1γ: n = 3; GFP-HDAC5: n = 7; GFP: n = 3; statistical analysis by Tukey-test).

Using this system, we first used time-lapse microscopy to measure the localization dynamics of GFP-HP1α and GFP-HDAC5 during transient expression, and their connection to gene expression in a cell population stably expressing the rTetR-antiGFP nanobody fusion and the TagRFP reporter. We observed that both GFP-tagged HP1α and HDAC5 mediated fast silencing of the reporter, as indicated by the decrease in TagRFP signal as early as 24 hours after dox addition (Fig. 1b). Furthermore, we found that while HP1α localizes mostly in the nucleus (Fig. S1a), HDAC5 was seen translocating in and out of the nucleus (Fig. S1b), as previously reported^29^. In cells where there was dilution of the GFP expression vector and thus the expression of the CR was lost, the TagRFP reporter reactivated after ~50 hours (Fig. 1b). At the end of five days of recruitment (dox treatment), the TagRFP reporter was still silent in cells that were still expressing GFP (Fig. 1c).

Based on these promising results, we then used flow cytometry to measure reporter silencing after 5 days of recruitment in cells ectopically expressing GFP-HP1α, GFP-HP1β, GFP-HP1γ, or GFP-HDAC5. By the end of five days of transient expression and recruitment, all CRs tested were able to silence the reporter in a dox-dependent manner in the majority of the cells still expressing the GFP-tagged CR (Fig. 1d). In addition, recruitment of GFP alone does not lead to silencing of the reporter, which suggests the silencing observed is dependent on the activity of the recruited CR (Fig. 1d). HP1α, HP1β, and HDAC5 lead to a higher fraction of cells silenced compared to HP1γ, consistent with their reported roles in silencing and association with heterochromatin^30,31^. Surprisingly, HP1γ, which associates with actively transcribed regions and plays a role in transcriptional elongation via RNA Polymerase II^32^, still leads to silencing in a majority of cells (Fig. 1d). Increasing the number of antiGFP nanobodies fused to a single rTetR to 8 did not increase the fraction of cells silenced (Fig. S1c), suggesting that a single nanobody is sufficient for silencing in our reporter system.

### Nanobodies against DNMT1 and HP1 can silence a reporter gene and confer epigenetic memory

Encouraged by the results with the antiGFP nanobody, we proceeded to test two existing nanobodies against endogenous chromatin regulators, antiHP1^33^ and antiDNMT^34^, for their capacity to silence and induce epigenetic memory. The antiHP1 nanobody was shown to bind to all three isoforms of HP1^33^, while the antiDNMT1 nanobody has previously been used to immunoprecipitate endogenous DNMT1 from whole cell lysate^34^. Using a similar system as the antiGFP nanobody, we cloned fusions between rTetR and either antiHP1 or antiDNMT1, and stably integrated them into HEK293T cells containing the TagRFP reporter with TetO sites at the AAVS1 locus (Fig. 2a). Both nanobodies mediated dox dependent silencing upon recruitment at the reporter for 5 days, albeit weaker than KRAB (Fig. 2b), a known strong repressor^35^ used in CRISPRi^1^. While KRAB can strongly silence nearly all (97.9%) of the cells, antiHP1 and antiDNMT1 mediated silencing were weaker resulting in 16.8% and 28.5% of cells silenced, respectively (Fig 2b). Surprisingly, silencing by antiHP1 is weaker than GFP-tagged HP1 proteins when mediated by the antiGFP nanobody (16.8% vs. 64% to 87% depending on the HP1 isoform recruited, Fig. 1d). This difference in silencing may be due to the nanobody’s weak binding affinity to HP1 or due to low levels of endogenous HP1 protein in the cell. In addition, treatment of the cells with 5-aza-2’-deoxycytidine (5-aza-2’), a DNA methyltransferase inhibitor, during recruitment abolished antiDNMT1 mediated silencing but not KRAB or antiHP1 (Fig. S2a), indicating that DNA methyltransferases are required for antiDNMT1 mediated silencing. On the other hand, treatment with chaetocin, a broad-spectrum inhibitor of lysine histone methyltransferases, reduced all three effectors’ ability to silence (Fig. S2b), suggesting that histone methylation is involved in both antiHP1 and antiDNMT1 mediated silencing.

**Figure 2:**
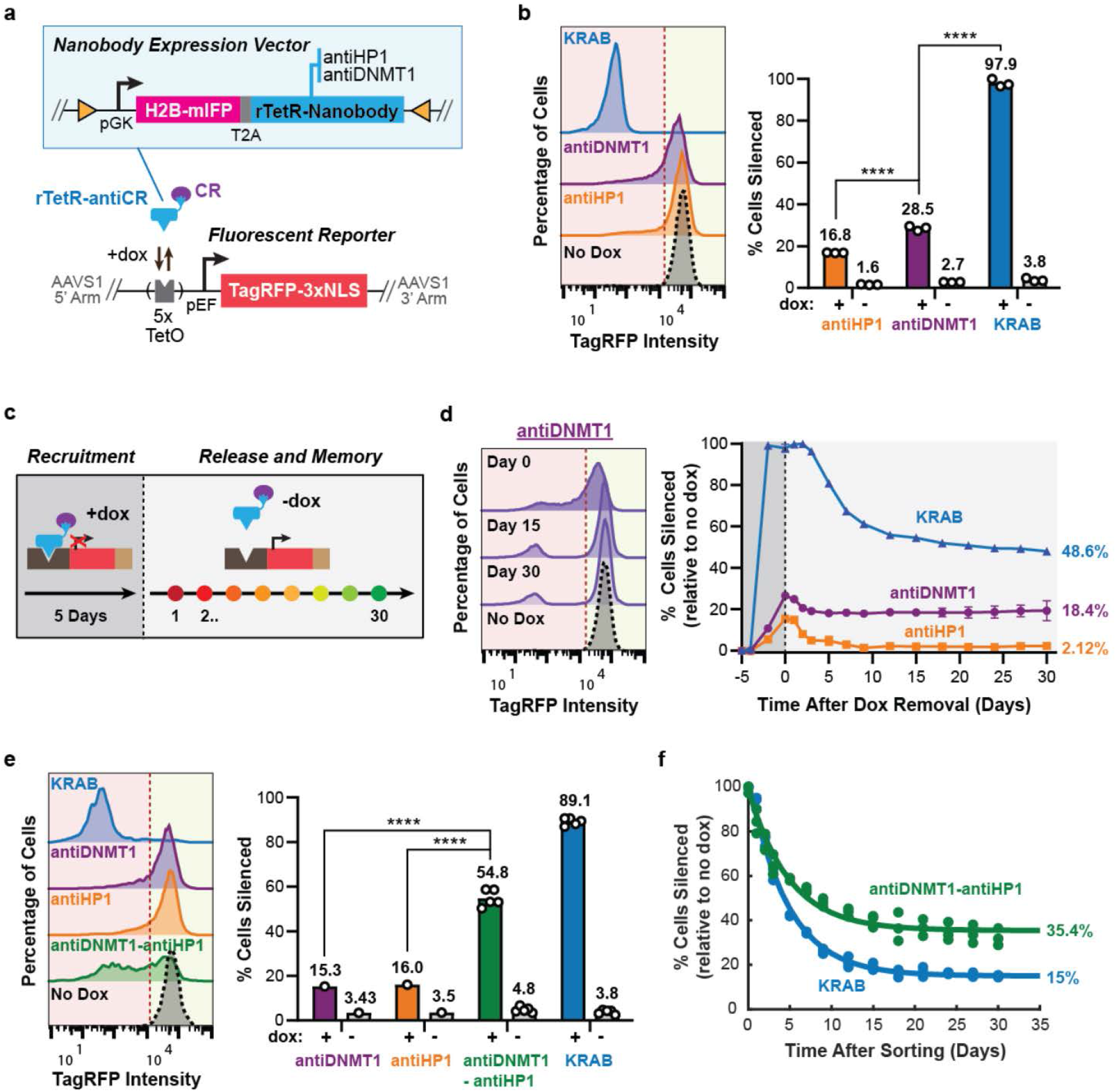
Nanobody-mediated recruitment of endogenous chromatin regulators can silence gene expression and confer memory. (**a**) Construct for constitutive coexpression of H2B-mIFP and nanobody against CR (antiHP1 or antiDNMT1) fused with the rTetR DNA binding protein (blue box, top) were expressed in HEK293T cells containing a TagRFP reporter (bottom). (**b**) (left) Fluorescence distributions of the TagRFP reporter after 5 days of recruitment (dox treatment) in cells stably expressing the nanobody constructs or rTetR-KRAB were analyzed by flow cytometry to determine the percentage of cells with the reporter silenced (left of the red dotted line). (right) Means of cells silent were calculated from 3 replicates (statistical analysis by Tukey-test). (**c**) Experimental design for investigating epigenetic memory: rTetR-effectors were recruited to the reporter for 5 days (+dox) and then released (−dox). Memory was monitored after dox removal via flow cytometry throughout 30 days. (**d**) Silencing and memory dynamics data (right) for the experiment described in (c) with representative flow cytometry histogram for antiDNMT1 at day 0, 15, 30 after dox removal (left). The percentage of cells silenced was normalized to the no dox control to adjust for any background silencing (Methods). Means are from 3 replicates. Standard deviations are plotted but are too small to show for most data points. (**e**) (left) Reporter fluorescent distributions and percent cells silent (right) after transient expression of rTetR-effector fusions and 5 days of dox treatment. Each dot is an independently transfected biological replicate (antiDNMT1: n = 1; antiHP1: n = 1; antiDNMT1-antiHP1: n = 5; KRAB: n = 5; statistical analysis by one-sample *t*-test). (**f**) Cells silenced by KRAB and antiDNMT1-antiHP1 in (e) were sorted after 5 days of dox treatment and analyzed by flow cytometry for memory. Each time point contains 3 biological replicates (individual dots). Data were fitted with an exponential decay curve (lines, Methods).

After recruiting the effectors for 5 days with dox, we released the cells from dox and assessed epigenetic memory by measuring the percentage of cells still silenced over 30 days with flow cytometry (Fig. 2c). We found that silencing mediated by antiHP1 was almost completely reversible (Fig. 2d; orange line), which is consistent with previous reports that silencing by HP1 is associated with limited and partial memory, with the majority of the cells reactivating gene expression over time^30,36^. In contrast, KRAB and antiDNMT1 led to more permanent memory, with about 50% and 65% cells irreversibly silenced out of the total initially silenced, respectively (Fig. 2d; blue and purple line). These observations were consistent with the reactivation dynamics of KRAB in a different mammalian cell line (CHO cells)^37^ and the role of DNA methyltransferases in mediating epigenetic memory^5–7,38^. Overall, these results demonstrate the first use of a nanobody that can mediate gene silencing and impart epigenetic memory.

Having the option to transiently express these nanobodies and bypass the need to stably integrate them in cells may provide greater flexibility for transcriptional control. Therefore, we tested how effective these nanobodies were at silencing when transiently expressed in our TagRFP reporter cell line. AntiHP1 and antiDNMT1 led to weak silencing of the reporter after 5 days of dox treatment (Fig. 2e), similarly to the stable expression results. In addition to testing the nanobodies individually, we fused the two nanobodies together with a short flexible linker to see if this would enhance gene silencing and memory at the reporter gene (Fig. S2c). Transient expression and recruitment of rTetR-antiDNMT1-antiHP1 led to about 55% of the cells being silenced which is more than each nanobody individually (Fig. 2e). In order to measure epigenetic memory independent of silencing efficiency, we decided to sort the silenced (TagRFP negative) cells at the end of 5 days of dox treatment and measure their persistence of silencing for 30 days. We found that the antiDNMT1-antiHP1 fusion had improved epigenetic memory over KRAB, with 35.4% cells still silent at 30 days post sorting versus 15% cells, respectively (Fig. 2f). These results show for the first time that combining two nanobodies that bind different CRs can be used to enhance gene silencing as well as epigenetic memory, and demonstrates the promising prospect of using nanobodies in combination with other CRs to improve transcriptional control.

### Recruitment of antiDNMT1 improves silencing speed and epigenetic memory of other CRs at the reporter

Realizing the potential of using nanobodies in combination with other CRs, we decided to test the antiDNMT1 nanobody in combination with KRAB (Fig. S3a). Since KRAB has strong, fast silencing and antiDNMT1 has long-lived epigenetic memory, we hypothesized that combining KRAB and antiDNMT1 might allow us to build a small tool for fast silencing and long-lived memory that can be delivered transiently. Transient expression and recruitment of rTetR-KRAB-antiDNMT1 to the reporter for 5 days resulted in strong silencing (87.8%), on the same level as KRAB alone (88.9%) (Fig. 3a), but demonstrated improved memory (Fig. 3b; pink vs. blue line). This increase is especially important for experiments that involve transient KRAB expression, as epigenetic memory of KRAB when transiently expressed (Fig. 3b; blue line, 15%) is lower than in cell lines stably expressing KRAB (Fig. 2d; blue line, 48.6%), presumably due to the dilution of the plasmid. The further addition of the antiHP1 nanobody to the KRAB-antiDNMT1 fusion did not increase silencing or memory further (Fig. 3a-b; brown). Moreover, when co-recruiting KRAB and antiDNMT1 simultaneously at the reporter via separate fusions to rTetR, epigenetic memory was even lower than KRAB recruitment alone (Fig. S3b). We measured a similar decrease in memory and silencing upon co-recruitment of separate fusions of rTetR with antiHP1 and antiDNMT1 (Fig. S3b and Fig 2e). These observed reductions in epigenetic memory and silencing may be due to effector binding competition at the five TetO sites and suggest that direct fusions of multiple CRs might be preferable to independent co-recruitment at a locus for gene transcriptional control.

**Figure 3:**
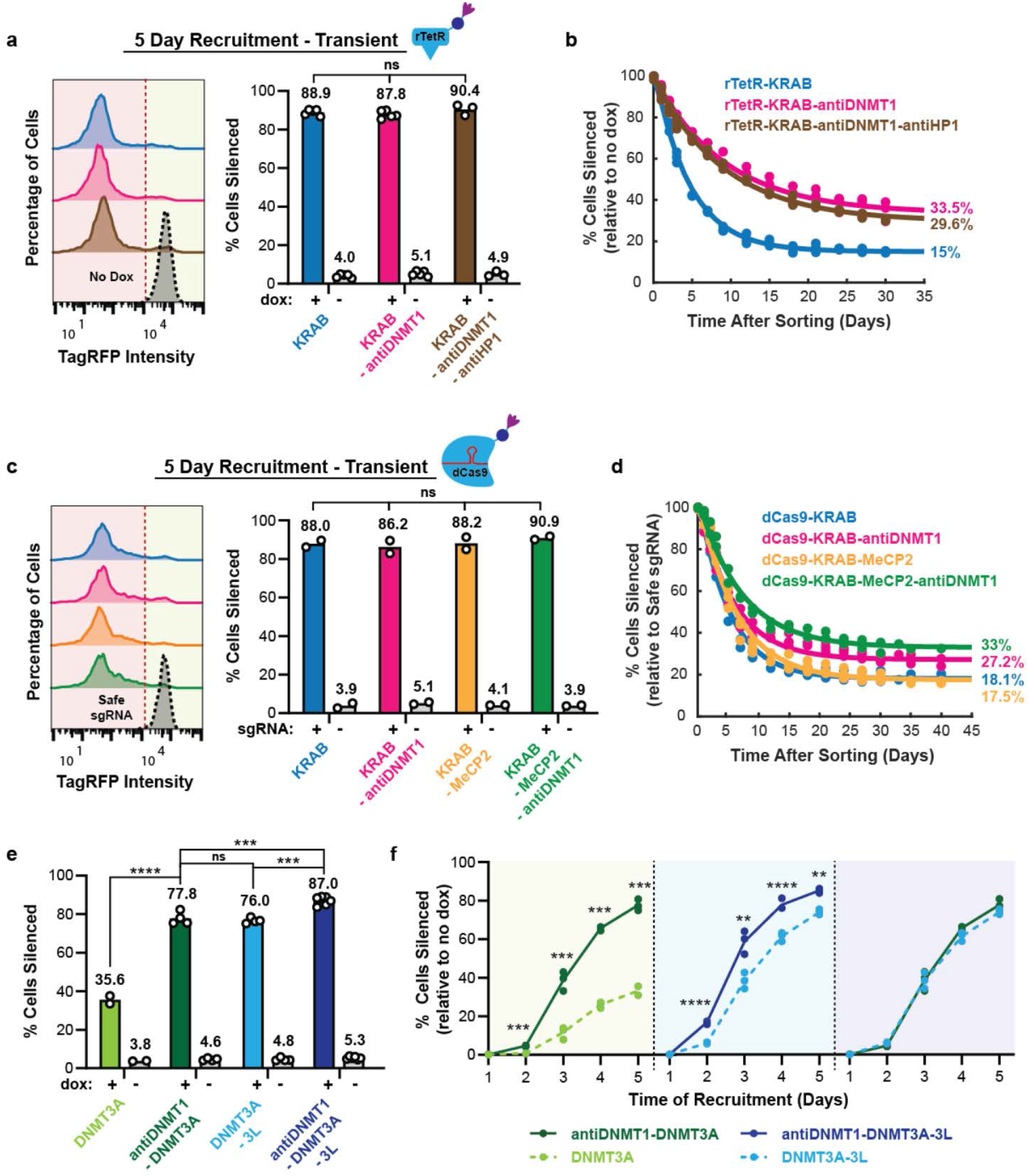
Nanobody-mediated enhancement of KRAB and DNMT3A repression. **(a)** (left) Fluorescence distributions of the reporter gene after transient expression of rTetR-effector fusions and recruitment by dox treatment for 5 days, measured using flow cytometry. (right) The means of percent cells silent (to the left of the red dotted line in the left-side plots) are shown as bars. Each dot represents an independently transfected biological replicate (KRAB: n = 5; KRAB-antiDNMT1: n = 5; KRAB-antiDNMT1-antiHP1: n = 3; statistical analysis by one-way ANOVA). **(b)** After recruitment with rTetR-effector fusions for 5 days, silenced cells were sorted, and memory dynamics were measured by flow cytometry throughout 30 days. Data represent 3 biological replicates and are fitted with an exponential decay curve (Methods). (**c**) Fluorescence distributions after transient expression and targeting of dCas9-effector fusions to the TetO sites upstream of the reporter gene (+) or to a safe-targeting control site (−) for 5 days. Means are from 2 biological replicates (statistical analysis by one-way ANOVA). (**d**) After targeting the dCas9-effector fusions for 5 days, silenced cells were sorted, and memory dynamics was measured by flow cytometry throughout 40 days. Each dot is a biological replicate (KRAB: n = 2; KRAB-antiDNMT1: n = 3; KRAB-MeCP2: n = 3; KRAB-MeCP2-antiDNMT1: n = 2) and are fitted with an exponential decay curve (Methods). (**e**) Transient expression and recruitment of rTetR-based DNA methyltransferase combinations to the reporter gene for 5 days (DNMT3A: n = 2; antiDNMT1-DNMT3A: n = 4; DNMT3A-3L: n = 4; antiDNMT1-DNMT3A-3L: n = 6; statistical analysis by Tukey-test). (**f**) Corresponding silencing dynamics of rTetR-based combinations in (e) throughout 5 days of recruitment (n = 3 replicates; statistical analysis by Tukey-test).

To expand the use of this tool for endogenous gene silencing, we tested the KRAB-antiDNMT1 fusion with the dCas9 system at the reporter gene (Fig. S3c). Single guide RNAs (sgRNAs) were designed to target the 5x TetO binding site upstream of the reporter and a genomic site with no annotated function to serve as a negative control (called safe-targeting guide). When the programmable dCas9-KRAB-antiDNMT1 was targeted to the reporter gene for 5 days, there was strong silencing and improvement in memory similar to rTetR (Fig. 3c-d; 27.2% vs. 18.1%). Recruitment of dCas9-KRAB-antiDNMT1 not only demonstrated improved memory over dCas9-KRAB but also over a newly developed combined repressor, dCas9-KRAB-MeCP2^4^, (Fig. 3d; 27.2% vs. 17.5%). In our system, KRAB-MeCP2 has the same memory as KRAB alone (Fig. 3d; 18.1% vs. 17.5%) and the addition of antiDNMT1 to this fusion resulted in a similar improvement in memory as when added to KRAB (Fig. 3d; 33% for KRAB-MeCP2-antiDNMT1 vs. 27.2% for KRAB-antiDNMT1). These results suggest that the antiDNMT1 nanobody can be used to enhance the permanent silencing ability of CRs. However, when we targeted the dCas9-KRAB-antiDNMT1 fusion to the *CXCR4* endogenous gene, we did not observe an increase of epigenetic memory compared to dCas9-KRAB alone (Fig. S3d-e), suggesting that the KRAB-antiDNMT1 tool requires further systematic characterization with respect to genomic locus and promoter type. While the level of memory seen after rTetR-KRAB-antiDNMT1 recruitment at the reporter is smaller than previously observed with the triple combination KRAB-DNMT3A-Dnmt3L^5,7^, KRAB-antiDNMT1 is about three times smaller (Fig. S3a; ~580 bp vs. ~1770 bp) and thus may be a more appealing tool for viral-based methods.

Recruitment of the catalytic domain of DNMT3A at a gene locus can induce DNA methylation and stable gene repression^39,40^. However, because DNMT3A alone typically leads to slow transcriptional repression, it is common to combine it with other CRs to enhance its effects^5–7^. Realizing the potential of the antiDNMT1 nanobody in improving the gene repressive effects of KRAB, we wanted to test if combining the nanobody with the catalytic domain of DNMT3A would enhance it as well. When the rTetR-antiDNMT1-DNMT3A fusion was recruited to the reporter gene via rTetR for 5 days, it led to stronger and faster silencing when compared to DNMT3A alone (Fig. 3e-f; dark green vs. light green). In addition, it has been shown that DNMT3L can enhance the catalytic activity of DNMT3A^41–43^. Consistent with previous work, the addition of the C-terminal domain of mouse Dnmt3L and the catalytic domain of DNMT3A enhanced silencing of our reporter from 35.6 to 76 percent (Fig. 3e; light blue). Surprisingly, the addition of the smaller antiDNMT1 nanobody to DNMT3A led to a similar improvement in silencing as the larger Dnmt3L domain (Fig. 3e-f; dark green vs. light blue). The antiDNMT1 nanobody further improved silencing when added to the DNMT3A-3L fusion (Fig. 3e-f; dark blue vs. light blue). In fact, of the different rTetR fusion combinations tested, the antiDNMT1-DNMT3A-3L triple fusion was by far the strongest (Fig. 3e; dark blue) resulting in about 87% of the cells being silenced at 5 days of dox. In summary, the antiDNMT1 nanobody improved the speed of silencing in all combinations with DNMT3A (Fig. 3f). All fusions containing rTetR-DNMT3A, including the ones containing antiDNMT1, led to permanent epigenetic memory at our reporter gene (Fig. S4a). We also see a similar increase in the speed of silencing of the reporter gene when we fused antiDNMT1 to the histone deacetylase enzyme HDAC4 (Fig. S4b). These promising results suggest the fusion of a small antiDNMT1 nanobody to CRs may serve as a new way to enhance silencing or memory.

### Nanobody-mediated recruitment of chromatin regulators for synthetic circuit control

These nanobody-based tools for controlling gene expression and epigenetic memory could serve as devices in synthetic circuits for detecting and recording signals. Cellular stopwatches and recording devices are important components of synthetic biology circuits^44^. The response of the antiDNMT1 nanobody presents a unique opportunity of implementing a very compact stopwatch that can both measure and record the duration of a signal. The desired signal can be coupled to the expression of rTetR-antiDNMT1 that in turn can be recruited upstream of an output gene encoding for fluorescence, signaling molecules, or proteins involved in cell death or survival (Fig. 4a). The addition of dox starts the time recording session, while removal of dox ends it. Since silencing by antiDNMT1 recruitment is quite slow, it leads to a linear response in the percentage of cells silent as a function of signal duration (Fig. 4b). Moreover, since antiDNMT1 recruitment leads to permanent memory in the majority of cells silenced, the signal duration can be recorded as a percentage of cells in the population that have the output gene off (Fig. 4c). Coupling antiDNMT1 expression or recruitment to another cellular signal (such as a cytokine or a hormone) would allow one to measure and permanently record the total duration of that signal as the fraction of cells with the reporter silenced.

**Figure 4.**
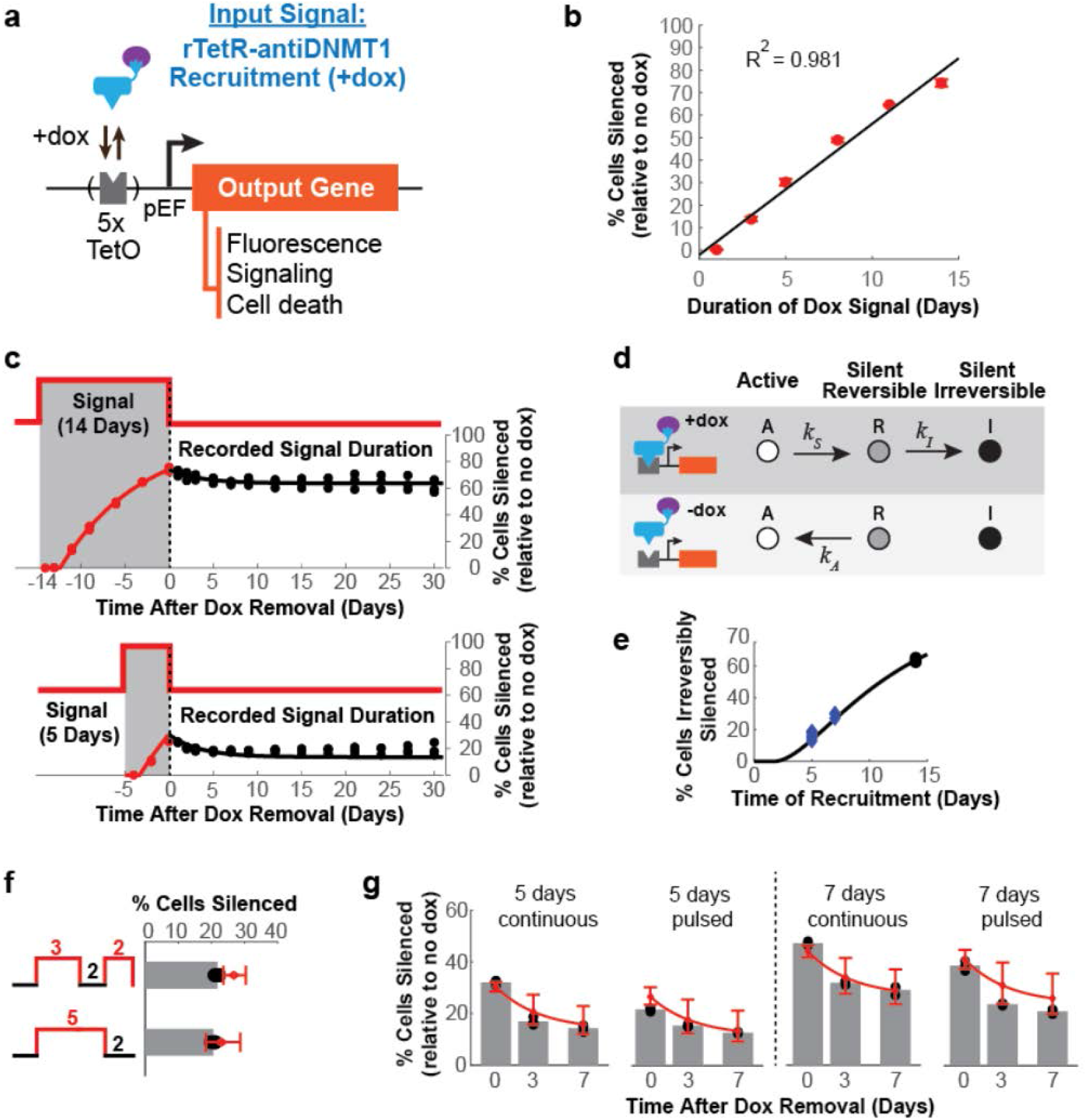
Nanobodies as signal detection and recording tools. (**a**) Schematic of a device for measuring and recording signal duration. The input signal is coupled to the recruitment of rTetR-antiDNMT1 near the pEF promoter to silence an output gene. (**b**) Percentages of cells with TagRFP reporter silenced as measured by flow cytometry at the end of the indicated dox signal durations in a cell line stably expressing rTetR-antiDNMT1. Means and standard deviations of experimental data from 3 replicates (red dots) and linear fit (black, Methods). (**c**) Percentage of cells with TagRFP silenced after different signal (dox treatment) durations: 14 days (top) and 5 days (bottom). The gray shaded regions (negative numbers) indicate the period with dox. Dashed lines indicate dox removal. Continuous lines represent fits to the model in (d) for the 14 days data, and predictions of the model for the 5 day data (Methods, Supplementary Text). (**d**) Three-state model of silencing by antiDNMT1 during recruitment (+dox, top) and during release (−dox, bottom). (**e**) Percentage of cells irreversibly silent after different durations of recruitment (dox treatment) predicted by the model in (d) are plotted as a black line (Methods). Experimental data recorded at 7 days during the release period shown as black dots if used for model fitting (14 days), or blue diamonds if not used in the fit. (**f**) Percentages of cells silenced relative to no dox controls for pulsed recruitment (top: 3 days +dox, 2 days −dox, 2 days +dox) compared to continuous recruitment for the same duration (bottom: 5 days +dox, 2 days −dox). Experimental data from 3 replicates shown as black dots, means as gray bars, and model predictions with 95% CI in red (Methods). (**g**) Percentages of cells silenced relative to no dox controls for pulsed recruitment versus continuous recruitment, recorded at the same time after dox removal, plotted as in (f).

The linear response to signal duration and its recording in the fraction of irreversibly silent cells can be described using a 3-state model of gene control (Fig. 4d, Methods)^37^. In this model, active cells (A) silence at a slow rate (*k*_*S*_) during recruitment of antiDNMT1. They first reach a reversible silent state (R), and can transition from this to an irreversibly silent state (I) with a rate *k*_*I*_. After release of the nanobody, the reversibly silent population reactivates at a rate *k*_*A*_, while the irreversibly silent cells remain silent. By fitting the experimental data for antiDNMT1 silencing and reactivation of the 14-day time-course in Fig. 4c (Methods), we can extract the three rates and their 95% confidence intervals: *k*_*S*_ = 0.11/day (0.1 - 0.12), *k*_*A*_ = 0.29/day (0.12 - 0.45), *k*_*I*_ = 0.37/day (0.31 - 0.43). Because the exponential rate of silencing is quite low, it results in an approximately linear fraction of cells silenced as the duration of signal (dox recruitment) increases (Fig. 4b-c, red dots). Similarly, the fraction of irreversibly silent cells that record signal duration looks approximately linear with signal duration for a large range of signals (Fig. 4e).

Once calibrated, the 3-state phenomenological model can be used to predict the fraction of cells silent over time for new types of signals. For example, the model predicts that the fraction of cells silenced at the end of a 5 day pulsed signal (3 days of dox, 2 days of no dox, and then 2 days of dox) is approximately the same as at the end of a continuous 5 day signal (5 days of dox and then 2 days of no dox), matching experimental data (Fig. 4f). This model also predicts that continuous signals result in a small but systematically higher level of cells permanently silenced compared to interrupted signals of the same total duration, which is also consistent with experimental data (Fig. 4g). Ultimately, this type of modelling can be used to program the fraction of cells active or silenced for synthetic biology applications.

## Discussion

In this study, we developed a technology that uses nanobodies to recruit endogenous CRs at a target gene for transcriptional and epigenetic memory control in mammalian cells. We showed that the antiGFP nanobody that binds to GFP-tagged chromatin regulators (GFP-CR) can mediate gene silencing when recruited to a reporter gene. Nanobodies against endogenous chromatin regulators (antiHP1 and antiDNMT1) can mediate silencing as well. Moreover, these nanobodies led to stronger silencing or more persistent epigenetic memory when combined with each other or with other CRs (KRAB, DNMT3A, and HDAC4). These results highlight the use of nanobodies as a method to enhance gene silencing by combining them with other effectors and open new avenues for development, characterization, and application of compact nanobody-based tools in gene regulation.

While here we used the antiGFP nanobody mainly for an initial proof of concept experiment, to show that targeted recruitment of nanobodies against GFP-CR fusions can mediate gene silencing, this approach can be further extended to elucidate how chromatin regulator localization and gene expression are interconnected. Different chromatin regulators are known to mediate liquid-liquid phase separation (LLPS) and translocate in and out of the nucleus, and the dynamics of these localization are thought to regulate gene expression. For example, HP1 localizes primarily to the nucleus, can form phase separated regions that partition heterochromatin^45,46^, and mediate silencing and partial epigenetic memory when recruited at a gene^30^. Class IIa HDACs (4, 5, 7 and 9) translocate in and out of the nucleus in a cell-cycle dependent manner, and this shuttling phenotype is believed to spatially control their ability to interact with repressive complexes in the nucleus and thus their ability to repress transcription^47–49^. Other localization patterns have been observed for other HDACs as well, ranging from being mostly in one subcellular compartment (nucleus or cytoplasm) to localizing to either compartment depending on the cell type^29^. Our nanobody-based recruitment tool has the potential to be applied to all HDACs (11 total) to better help understand their localization pattern and role in gene regulation changes in different cell types. However, in order to connect the fast changes in CR localization (shorter than a cell cycle) to gene expression, one needs to use an mRNA-based reporter (e.g. MS2-MCP or PP7-PCP systems), rather than a protein reporter (e.g. mCitrine or TagRFP).

Gene regulation using the antiGFP nanobody can also be used in conjunction with recently engineered cell line collections containing a GFP knock-in at loci expressing specific chromatin regulators and transcription factors^50,51^. These cell lines express CR-GFP fusions at endogenous levels, and thus allow characterization of localization, dynamics of phase separation properties, and gene regulation via the antiGFP nanobody at concentrations that are biologically relevant, without overexpression of the regulators. Moreover, some of these cell collections consist of stem cells, either human induced pluripotent cells (hiPSCs)^50^ or mouse embryonic stem cells (mESCs)51, that can be differentiated to different cell types. Therefore, introducing the reporter-antiGFP system in these lines would allow simultaneous characterization of CR and gene regulation dynamics in various differentiated cell types that have the same genetic background without the need for additional cell engineering. Moreover, development of nanobodies against smaller tags, for example against the small C-terminal of the split GFP, GFP11, would make this approach easier, as knock-in of GFP11 into endogenous loci can be done in a higher throughput manner^52^.

We also show that nanobodies can be used in synthetic circuits for detecting and recording signals. Using experimental data for antiDNMT1 mediated silencing and reactivation at the TagRFP reporter, we were able to derive a 3-state model of gene control for this nanobody that can be used to predict the fraction of cells silent over time for new types of signals. This interplay between experiment and mathematical modeling is key to informing the design and fine-tuning of synthetic circuits, which can then be applied to engineer larger, more complex networks. In addition, models that accurately predict the behavior of a system can help us engineer new cellular outputs without having to perform large numbers of trial-and-error experiments. The framework we developed for antiDNMT1 can be expanded to study the capabilities of other nanobodies and used to optimize gene circuits that take advantage of the specific nanobodies silencing and memory characteristics.

Development of nanobodies against other chromatin regulators and transcription factors besides antiHP1 or antiDNMT1 could lead to even more versatile gene regulation and epigenetic control. For example, recruitment of *de novo* DNA methylases DNMT3A and DNMT3B enables DNA methylation of promoters, enhancers, and other regulatory regions (e.g. CTCF sites) for transcriptional repression^17,39,40,53^. Nanobodies designed against these de novo methylases could be used for locus specific methylation in the same manner, and possibly avoid off-target methylation toxicity associated with overexpression of the methylases^17,54^. Another class of chromatin regulator are those associated with euchromatin and gene activation: histone acetylases (p300)^9^, DNA hydroxymethylases that mediate removal of DNA methylation (Tet1-3)^39,55,56^, and ATP-dependent chromatin remodeling complexes (BAF)^36^. Nanobodies against these complexes could be used for activation of heterochromatic loci. When developing new nanobodies for gene control it is important to target CRs that are abundant in the cells of interest and to select nanobodies with high binding affinity to their target CRs. Development of such functional nanobodies that can properly fold in the cellular environment and bind to different chromatin complexes with high affinity without disrupting their function requires new high-throughput screening methods inside human cells. The reporter system and methods described here could serve as a platform for such nanobody functional screens.

In order to extend the use of nanobodies beyond control of reporter genes, it is necessary to take a systematic high-throughput approach that tests variability with target locus and cell type. Variability in the effect on gene expression is not specific to nanobody-mediated control, but has been reported for direct fusions of dCas9 and full-length chromatin regulators^5,6^. For example, persistent epigenetic memory requires both DNA methylation by DNMT3A-3L and histone methylation by EZH2 (H3K27me3) or KRAB (H3K9me3)^6^. However, which histone methylase is necessary for memory depends on the specific gene targeted^5,6^. Even at genes where KRAB and DNMT3A-DNMT3L co-recruitment mediates persistent memory in one cell type, the combination sometimes fails to impart memory in a different cell type^5^. These results suggest we also need a clearer understanding of the co-factors each CR interacts with, and how their expression and activity varies with genomic locus and cell type. Moreover, the off-target effects of different tools can also vary for different dCas9 fusions^57^, and thus would have to be systematically measured. Compact tools such as nanobodies, that can be more easily cloned and delivered alone and in different combinations, would facilitate systematic measurements of on target variability and off target effects of epigenetic editing tools.

## Supporting information

Van2020 Supplement

## Acknowledgements

This work was supported by the NIH T32 Training Grant T32GM007276 (M.V.V.) and NIH MIRA R35GM128947-01 (L.B.), and a BWF-CASI Award (L.B.). We thank Frank Perez and Sandrine Moutel (Institut Curie) for the antiHP1 nanobody plasmid, Rajarshi Ghosh and Jan Liphardt (Stanford University) for the antiGFP nanobody plasmid, Stanley Qi (Stanford University) and members of his lab for allowing us to use their flow cytometer, Josh Tycko and Joydeb Sinha (Stanford University) for providing us with plasmids, and members of the Bintu Lab for useful feedback on the manuscript.

## Author Contributions

M.V.V. and L.B. designed the study. M.V.V. generated the DNA constructs and cell lines, performed the flow cytometry experiments and associated data analysis. T.F. performed the experiments and data analysis for the time-lapse movies. L.B. derived and fit the 3-state gene control model. M.V.V. and L.B. wrote the manuscript with inputs from T.F..

## Competing Interests

The authors declare no competing interests.

## Methods

### Plasmid construction

The TagRFP reporter (5xTetO-pEF-TagRFP-3xNLS) construct was assembled using a AAV zinc finger donor vector backbone (Addgene #22212) containing a promoter-less splice-acceptor (SA) upstream of a puromycin resistance gene and homology arms against the *AAVS1* locus. Three elements of the reporter were amplified from the following sources: five TetO binding sites upstream of a pEF promoter from PhiC31-Neo-ins-5xTetO-pEF-H2B-Citrine-ins (Addgene #78099), TagRFP-T from pEN_ERK.KTR-tagRFP-T, and 3xNLS from pEN_mCherry-NLS (both gifts from Joydeb Sinha & Mary Teruel, Stanford). These components were cloned into the AAV donor vector backbone using Gibson Assembly.

The plasmids containing the rTetR-effector fusions were cloned into the PB-CMV-MCS-EF1α-Puro PiggyBac vector backbone (System Biosciences #PB510B-1), which was further modified via Gibson Assembly with the following components: PGK promoter from pSLQ2818, mIFP from pSLQ2837-1 (both gifts from Tony Gao & Stanley Qi, Stanford)^58^, H2B-rTetR-Zeo from pEx1-pEF-H2B-mCherry-T2A-rTetR-KRAB-Zeo (Addgene #78352). Each effector was PCR amplified from the following plasmid: antiGFP nanobody^59^ - gift from Rajarshi Ghosh & Jan Liphardt, Stanford; antiDNMT1 nanobody - purchased from ChromoTek; antiHP1 nanobody^33^ - gift from Sandrine Moutel & Franck Perez, Institut Curie; KRAB - Addgene #84241; MeCP2 - Addgene #110821; and DNMT3A-3L - Addgene #71827.

Plasmids containing the dCas9-effector fusions were derived from the dCas9-KRAB vector backbone (Addgene #110820) and modified by Gibson Assembly with their respective effectors from sources listed above. The dCas9-effector fusions containing KRAB or KRAB-antiDNMT1 were further modified with mCitrine-NLS upstream of the dCas9 to allow for cell sorting and analysis of endogenous gene silencing. The sgRNA cloning vector (gift from Josh Tycko & Michael Bassik, Stanford, Addgene #89359) was modified to express mIFP or mCherry. Each sgRNA sequence (see Table S1) was cloned into the plasmid using the BlpI and BstXI cloning sites, as previously described^60^.

### Cell culture

Cells were cultured at 37°C under a humidified atmosphere with 5% CO2. HEK293T cells (Takara Bio #632180) were maintained in Dulbecco’s modified Eagle medium (DMEM; Gibco #10569010) supplemented with 25mM D-glucose (Gibco), 1 mM sodium pyruvate (Gibco), 1x GlutaMAX™ (Gibco), and 10% Tet Approved FBS (Clontech Laboratories). When cells reached 80% confluence, they were gently washed with 1x DPBS (Life Technologies) and passaged using 0.25% Trypsin (Life Technologies). For long-term storage, cells were resuspended in freezing media (10% DMSO (Sigma) and cell media) in a cryovial and frozen at −80°C.

### Stable cell lines construction

The reporter cell line was created by integrating the TagRFP fluorescent reporter at the first intron of the constitutively expressed gene *PPP1R12C* at the *AAVS1* locus in HEK293T cells. The integration of the reporter was performed by co-transfecting 1000 ng TagRFP reporter (5xTetO-pEF-TagRFP-3xNLS) donor plasmid and 500 ng of each TALEN arm (*AAVS1*-TALEN-L (Addgene #35431) targeting TGTCCCCTCCACCCCACA and *AAVS1*-TALEN-R (Addgene #35432) targeting TTTCTGTCACCAATCCTG). Cells were selected with 500 ng/mL puromycin (InvivoGen) starting 48 hours post-transfection for approximately 5 days or until all of the negative control cells died. Cells positive for TagRFP had two peaks representing the monoallelic and bi-allelic integration of the reporter at the *AAVS1* locus. Cells with the lower fluorescence peak (monoallelic) were sorted by fluorescence-activated cell sorting (FACS) using a Sony SH800 Cell Sorter with a 100um disposable chip. Each of the individual rTetR-effector plasmids was randomly integrated into this reporter line by co-transfecting 250 ng Super PiggyBac Transposase expression vector (System Biosciences #PB200PA-1) and 750 ng of rTetR-effector donor vector. These cells were selected with 60 µg/mL zeocin (InvivoGen) starting 48 hours post-transfection. All transfections were performed in 24-well plates using Lipofectamine 2000 (Invitrogen).

### Transient transfections

Approximately 70,000 cells were seeded per well in a 24-well plate, and the next day cells were transfected using Lipofectamine 2000 (Invitrogen) according to manufacturer instructions. For experiments in the antiGFP nanobody cell line, 1000 ng of each GFP-tagged chromatin regulator was delivered. For the transient silencing and reactivation experiments, 1000 ng of rTetR-effector expression vector was delivered to each well. 600 ng of dCas9-effector and 400 ng of sgRNA were co-delivered for the silencing and reactivation experiments involving dCas9 fusions. A maximum of 1000 ng of DNA vector was used per transfection.

### Acquisition of time-lapse movies

Approximately 200,000 reporter cells stably expressing rTetR-antiGFP were seeded per well in a 24-well plate, and the next day were transfected with 1000 ng GFP-CR plasmids using Lipofectamine LTX (Invitrogen), according to manufacturer’s instructions. We found that transfection with Lipofectamine LTX helps reduce cell death during time-lapse imaging. Six hours after transfection, cells were re-seeded at a density of approximately 20,000 cells in a 24-well imaging plate (ibidi #82406) coated with 2% Matrigel (Corning), left overnight in the incubator to adhere, and imaging was started early the next day. Imaging was done using a Leica DMi8 fluorescence microscope with Adaptive Focus Control (AFC), a 20X or 40X dry objective, and a Leica DFC9000 GT sCMOS camera. Fluorophores were excited using a Lumencor SOLA SE II light source. Images were automatically acquired every 15 minutes, using LAS X software (Leica Microsystems). Cells were grown in low-fluorescence imaging media, which consisted of FluoroBriteTM DMEM (Gibco) supplemented with 25mM D-glucose (Gibco), 1mM sodium pyruvate (Gibco), 1x GlutaMAX™ (Gibco), 10% Tet Approved FBS (Clontech Laboratories), and 1X Penicillin/Streptomycin (Gibco). The microscope was enclosed in an environmental control chamber (OkoLab) kept at 37°C and 5% CO2. Doxycycline (Tocris) was added to the imaging media to a final concentration of 1 μg/mL. The imaging media was changed daily for ~5 days (until the cells became too confluent to continue movies). Time-lapse movies were analyzed using ImageJ by visually tracking individual cell lineages and manually circling the area corresponding to the cell’s nuclei one hour after each cell division. Average fluorescence intensities of mIFP, TagRFP, and GFP within these contours of the cell nuclei were calculated and plotted based on their cell lineage using MATLAB (MathWorks).

### Gene expression analysis via flow cytometry

Cells expressing stably integrated or transiently transfected rTetR-effectors were assayed by flow cytometry during and after 5 days of 1 μg/mL doxycycline (Tocris) treatment. When indicated, cells were also treated with 1 μM 5-Aza-2’-deoxycytidine (Sigma) or 100 nM chaetocin (Cayman Chemical). Media containing small molecules were replaced daily. For experiments involving dCas9-effectors, cells were analyzed 5 days post-transfection and after being sorted for silencing (TagRFP negative cells). On the day of flow cytometry analysis, cells were harvested using 0.25% Trypsin (Life Technologies). A fraction of the cells (varying between one half to one twentieth, depending on cell density) were re-plated for the next time point. The remaining cells were resuspended in flow buffer (1x Hank’s Balanced Salt Solution (Life Technologies) and 2.5 mg/mL BSA (Sigma)) and filtered through a 40 μm strainer (Corning) to remove cell clumps. Cellular fluorescence distributions were measured with the CytoFLEX S Flow Cytometer (Beckman Coulter). The resulting data were analyzed with a custom MATLAB program called EasyFlow (https://antebilab.github.io/easyflow/). After cells were gated based on forward and side scatter, a manual gate was imposed on the TagRFP fluorescence to determine the percentage of silent cells for each sample. The gate was selected to contain 1-5% of the positive TagRFP signal in untreated cells. A total minimum of 20,000 events were recorded for each sample.

For experiments analyzing the expression of the endogenous *CXCR4* gene, cells were first trypsinized and then washed with 1% BSA (Sigma) in 1x DPBS (Life Technologies). Cells were then incubated on ice for 1 hour with monoclonal Brilliant Violet 421 (BV421) labeled anti-CXCR4 antibody (clone 12G5 (1:20); BioLegend). BV421 labeled IgG2a (clone MOPC-173 (1:20); Biolegend) served as a isotype control. Afterwards, cells were washed three times with 1% BSA/DPBS and then analyzed by flow cytometry for cells that were double positive for dCas9 (mCitrine) and sgRNA (mCherry).

### Statistical analysis

Data are displayed as individual points or as mean ± SD, with sample size indicated in the figure legend. Statistical significance was evaluated using GraphPad Prism 8. A one-way ANOVA with post hoc Tukey HSD (honestly significant difference) test or Student’s two-sample unpaired *t*-test were used for statistical analysis, with the exception of Fig. 2e (one-sample *t*-test). The one-sample *t*-test assumed that the hypothesized population mean for antiDNMT1 and antiHP1 were 15.3 and 16.0, respectively. Statistical significance of the data is indicated as follows: *p < 0.05; **p < 0.01; ***p < 0.001; ****p < 0.0001; ns = not significant.

### Gene control modeling and data fitting

The percentage of cells in each of the 3 states in the gene control model (Fig. 4d) was derived by solving the differential equations associated with the kinetic model (Supplementary Text), and all fits were done based on these equations, as follows:

In Fig. 2f, 3b, 3d, and S4a, the percentage of cells silent during the release period τ are fit to an exponential decay: *S*(τ) = (100 − *I*_1_)*e*^−*k*^*A*^τ^ + *I*_1_, where *I*_1_ is the percentage of cells irreversibly silenced at the end of the recruitment period, and *k*_*A*_ is the rate of reactivation.

For Fig. 4c, we fit the fraction of cells active (A = (1 - % cells silenced) / 100) after a recruitment signal (+dox) of duration *t* and reactivation period (−dox) of duration *τ*, to the following equation (derived in Supplementary Text, see equations (4)):

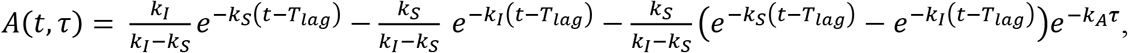

for all recruitment times *t* ≥ *T*_*lag*_, and *A*(*t*, τ) = 1 for *t* < *T*_*lag*_.

Note that for silencing time points, since τ = 0, the equation above simplifies to: *A*(*t*) = *e*^−*k*_*S*_^(*t*− *T*_*lag*_). All experimental data points from 3 independent replicates across 14-days of silencing (red points in Fig. 4c, top) and 30 days of reactivation (black points in Fig. 4c, top) were recorded in MATLAB as 3 vectors: *A* (percent of cells silent), *t* (time with dox) and τ (time after dox removal). For the silencing data points (+dox) *t* = 0,1, …, 14 and τ = 0, 0, …, 0. For reactivation data points (−dox) t= 14, 14, …, 14 and τ = 0, 1, 2, …, 30. These data were fit to the *A*(*t*, τ) equation above using MATLAB’s fit function with the NonlinearLeastSquares method, with *k*_*S*_, *k*_*I*_, *k*_*A*_, and *T*_*lag*_ as free parameters.

For pulsing experiments predictions in Fig. 4f&g, the full solutions *A*(*t*, τ), *R*(*t*, τ), and *I*(*t*, τ) were used to calculate the fractions of cells in each state for each period. For the first +dox period, the starting conditions were *A*_0_ = 1, *R*_0_ = 0, *I*_0_ = 0, and cells started silencing after a lag period *T*_*lag*_, so we used equations (4) in Supplementary Text. For the next periods, we applied the general equations (3) in Supplementary Text, using as starting conditions for each period the end points from the previous one. The *T*_*lag*_ period was only applied in the first dox induction period.

The linear fit in Fig. 4b was performed using MATLAB’s polyfit function with coefficient 1, and the goodness of fit was calculated using 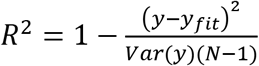, where *y* is the mean percent cells silenced at each time point from 3 replicates, and *N* is the number of time points.

#### Data Availability

All data and materials generated during this study are included in this manuscript and available from the corresponding author upon request.

## Notes

### Competing Interest Statement

The authors have declared no competing interest.

## References

1. Gilbert, L. a. et al. CRISPR-mediated modular RNA-guided regulation of transcription in eukaryotes. Cell 154, 442–451 (2013).

2. Schultz, D. C., Friedman, J. R. & Rauscher, F. J., 3rd. Targeting histone deacetylase complexes via KRAB-zinc finger proteins: the PHD and bromodomains of KAP-1 form a cooperative unit that recruits a novel isoform of the Mi-2alpha subunit of NuRD. Genes Dev. 15, 428–443 (2001).

3. Schultz, D. C., Ayyanathan, K., Negorev, D., Maul, G. G. & Rauscher, F. J., 3rd. SETDB1: a novel KAP-1-associated histone H3, lysine 9-specific methyltransferase that contributes to HP1-mediated silencing of euchromatic genes by KRAB zinc-finger proteins. Genes Dev. 16, 919–932 (2002).

4. Yeo, N. C. et al. An enhanced CRISPR repressor for targeted mammalian gene regulation. Nat. Methods 15, 611–616 (2018).

5. Amabile, A. et al. Inheritable Silencing of Endogenous Genes by Hit-and-Run Targeted Epigenetic Editing. Cell 167, 219–232.e14 (2016).

6. O’Geen, H. et al. Ezh2-dCas9 and KRAB-dCas9 enable engineering of epigenetic memory in a context-dependent manner. Epigenetics Chromatin 12, 26 (2019).

7. Mlambo, T. et al. Designer epigenome modifiers enable robust and sustained gene silencing in clinically relevant human cells. Nucleic Acids Res. 46, 4456–4468 (2018).

8. Kwon, D. Y., Zhao, Y.-T., Lamonica, J. M. & Zhou, Z. Locus-specific histone deacetylation using a synthetic CRISPR-Cas9-based HDAC. Nat. Commun. 8, 15315 (2017).

9. Hilton, I. B. et al. Epigenome editing by a CRISPR-Cas9-based acetyltransferase activates genes from promoters and enhancers. Nat. Biotechnol. 33, 510–517 (2015).

10. Cano-Rodriguez, D. et al. Writing of H3K4Me3 overcomes epigenetic silencing in a sustained but context-dependent manner. Nat. Commun. 7, 12284 (2016).

11. Kearns, N. A. et al. Functional annotation of native enhancers with a Cas9-histone demethylase fusion. Nat. Methods 12, 401–403 (2015).

12. Jacobs, S. A. et al. Specificity of the HP1 chromo domain for the methylated N-terminus of histone H3. EMBO J. 20, 5232–5241 (2001).

13. Platero, J. S., Hartnett, T. & Eissenberg, J. C. Functional analysis of the chromo domain of HP1. EMBO J. 14, 3977–3986 (1995).

14. Cowieson, N. P., Partridge, J. F., Allshire, R. C. & McLaughlin, P. J. Dimerisation of a chromo shadow domain and distinctions from the chromodomain as revealed by structural analysis. Curr. Biol. 10, 517–525 (2000).

15. Fuks, F., Hurd, P. J., Deplus, R. & Kouzarides, T. The DNA methyltransferases associate with HP1 and the SUV39H1 histone methyltransferase. Nucleic Acids Res. 31, 2305–2312 (2003).

16. Gowher, H. & Jeltsch, A. Molecular enzymology of the catalytic domains of the Dnmt3a and Dnmt3b DNA methyltransferases. J. Biol. Chem. 277, 20409–20414 (2002).

17. Lin, L. et al. Genome-wide determination of on-target and off-target characteristics for RNA-guided DNA methylation by dCas9 methyltransferases. Gigascience 7, 1–19 (2018).

18. Margot, J. B. et al. Structure and function of the mouse DNA methyltransferase gene: Dnmt1 shows a tripartite structure. J. Mol. Biol. 297, 293–300 (2000).

19. Ma, D., Peng, S., Huang, W., Cai, Z. & Xie, Z. Rational Design of Mini-Cas9 for Transcriptional Activation. ACS Synth. Biol. 7, 978–985 (2018).

20. Zetsche, B., Volz, S. E. & Zhang, F. A split-Cas9 architecture for inducible genome editing and transcription modulation. Nat. Biotechnol. 33, 139–142 (2015).

21. Truong, D.-J. J. et al. Development of an intein-mediated split-Cas9 system for gene therapy. Nucleic Acids Res. 43, 6450–6458 (2015).

22. Chew, W. L. et al. A multifunctional AAV-CRISPR-Cas9 and its host response. Nat. Methods 13, 868–874 (2016).

23. Adli, M. The CRISPR tool kit for genome editing and beyond. Nat. Commun. 9, 1911 (2018).

24. Hassanzadeh-Ghassabeh, G., Devoogdt, N., De Pauw, P., Vincke, C. & Muyldermans, S. Nanobodies and their potential applications. Nanomedicine 8, 1013–1026 (2013).

25. Beghein, E. & Gettemans, J. Nanobody Technology: A Versatile Toolkit for Microscopic Imaging, Protein–Protein Interaction Analysis, and Protein Function Exploration. Front. Immunol. 8, 446 (2017).

26. Rothbauer, U. et al. Targeting and tracing antigens in live cells with fluorescent nanobodies. Nat. Methods 3, 887–889 (2006).

27. Kirchhofer, A. et al. Modulation of protein properties in living cells using nanobodies. Nat. Struct. Mol. Biol. 17, 133–138 (2010).

28. Rothbauer, U. et al. A versatile nanotrap for biochemical and functional studies with fluorescent fusion proteins. Mol. Cell. Proteomics 7, 282–289 (2008).

29. Joshi, P. et al. The functional interactome landscape of the human histone deacetylase family. Mol. Syst. Biol. 9, 672 (2013).

30. Hathaway, N. A. et al. Dynamics and memory of heterochromatin in living cells. Cell 149, 1447–1460 (2012).

31. Zhang, C. L., McKinsey, T. A. & Olson, E. N. Association of class II histone deacetylases with heterochromatin protein 1: potential role for histone methylation in control of muscle differentiation. Mol. Cell. Biol. 22, 7302–7312 (2002).

32. Vakoc, C. R., Mandat, S. A., Olenchock, B. A. & Blobel, G. A. Histone H3 lysine 9 methylation and HP1gamma are associated with transcription elongation through mammalian chromatin. Mol. Cell 19, 381–391 (2005).

33. Moutel, S. et al. NaLi-H1: A universal synthetic library of humanized nanobodies providing highly functional antibodies and intrabodies. Elife 5, (2016).

34. Nishiyama, A. et al. Two distinct modes of DNMT1 recruitment ensure stable maintenance DNA methylation. Nat. Commun. 11, 1222 (2020).

35. Margolin, J. F. et al. Kruppel-associated boxes are potent transcriptional repression domains. Proc. Natl. Acad. Sci. U. S. A. 91, 4509–4513 (1994).

36. Braun, S. M. G. et al. Rapid and reversible epigenome editing by endogenous chromatin regulators. Nat. Commun. 8, 560 (2017).

37. Bintu, L. et al. Dynamics of epigenetic regulation at the single-cell level. Science 351, 720–724 (2016).

38. O’Geen, H. et al. dCas9-based epigenome editing suggests acquisition of histone methylation is not sufficient for target gene repression. Nucleic Acids Res. 45, 9901–9916 (2017).

39. Liu, X. S. et al. Editing DNA Methylation in the Mammalian Genome. Cell 167, 233–247.e17 (2016).

40. Vojta, A. et al. Repurposing the CRISPR-Cas9 system for targeted DNA methylation. Nucleic Acids Res. 44, 5615–5628 (2016).

41. Chedin, F., Lieber, M. R. & Hsieh, C.-L. The DNA methyltransferase-like protein DNMT3L stimulates de novo methylation by Dnmt3a. Proc. Natl. Acad. Sci. U. S. A. 99, 16916–16921 (2002).

42. Siddique, A. N. et al. Targeted methylation and gene silencing of VEGF-A in human cells by using a designed Dnmt3a-Dnmt3L single-chain fusion protein with increased DNA methylation activity. J. Mol. Biol. 425, 479–491 (2013).

43. Stepper, P. et al. Efficient targeted DNA methylation with chimeric dCas9-Dnmt3a-Dnmt3L methyltransferase. Nucleic Acids Res. 45, 1703–1713 (2017).

44. Dalchau, N. et al. Computing with biological switches and clocks. Nat. Comput. 17, 761–779 (2018).

45. Larson, A. G. et al. Liquid droplet formation by HP1α suggests a role for phase separation in heterochromatin. Nature (2017) doi:10.1038/nature22822.

46. Strom, A. R. et al. Phase separation drives heterochromatin domain formation. Nature 547, 241–245 (2017).

47. Grozinger, C. M. & Schreiber, S. L. Regulation of histone deacetylase 4 and 5 and transcriptional activity by 14-3-3-dependent cellular localization. Proc. Natl. Acad. Sci. U. S. A. 97, 7835–7840 (2000).

48. Wang, A. H. et al. Regulation of histone deacetylase 4 by binding of 14-3-3 proteins. Mol. Cell. Biol. 20, 6904–6912 (2000).

49. Kao, H. Y. et al. Mechanism for nucleocytoplasmic shuttling of histone deacetylase 7. J. Biol. Chem. 276, 47496–47507 (2001).

50. Roberts, B. et al. Systematic gene tagging using CRISPR/Cas9 in human stem cells to illuminate cell organization. Mol. Biol. Cell 28, 2854–2874 (2017).

51. Harikumar, A. et al. An Endogenously Tagged Fluorescent Fusion Protein Library in Mouse Embryonic Stem Cells. Stem Cell Reports 9, 1304–1314 (2017).

52. Leonetti, M. D., Sekine, S., Kamiyama, D., Weissman, J. S. & Huang, B. A scalable strategy for high-throughput GFP tagging of endogenous human proteins. Proceedings of the National Academy of Sciences 113, E3501–E3508 (2016).

53. McDonald, J. I. et al. Reprogrammable CRISPR/Cas9-based system for inducing site-specific DNA methylation. Biol. Open 5, 866–874 (2016).

54. Galonska, C. et al. Genome-wide tracking of dCas9-methyltransferase footprints. Nat. Commun. 9, 597 (2018).

55. Xu, X. et al. A CRISPR-based approach for targeted DNA demethylation. Cell Discov 2, 16009 (2016).

56. Choudhury, S. R., Cui, Y., Lubecka, K., Stefanska, B. & Irudayaraj, J. CRISPR-dCas9 mediated TET1 targeting for selective DNA demethylation at BRCA1 promoter. Oncotarget 7, 46545–46556 (2016).

57. Tycko, J. et al. Mitigation of off-target toxicity in CRISPR-Cas9 screens for essential non-coding elements. Nat. Commun. 10, 4063 (2019).

58. Gao, Y. et al. Complex transcriptional modulation with orthogonal and inducible dCas9 regulators. Nat. Methods 13, 1043–1049 (2016).

59. Ghosh, R. P. et al. A fluorogenic array for temporally unlimited single-molecule tracking. Nat. Chem. Biol. 15, 401–409 (2019).

60. Gilbert, L. A. et al. Genome-Scale CRISPR-Mediated Control of Gene Repression and Activation. Cell 159, 647–661 (2014).

