## Supplementary material for "Nanobody-mediated control of gene expression and epigenetic memory": Van2020 Supplement

Figure S1

Figure S2

Figure S3

Figure S4

Table S1

Supplementary Text

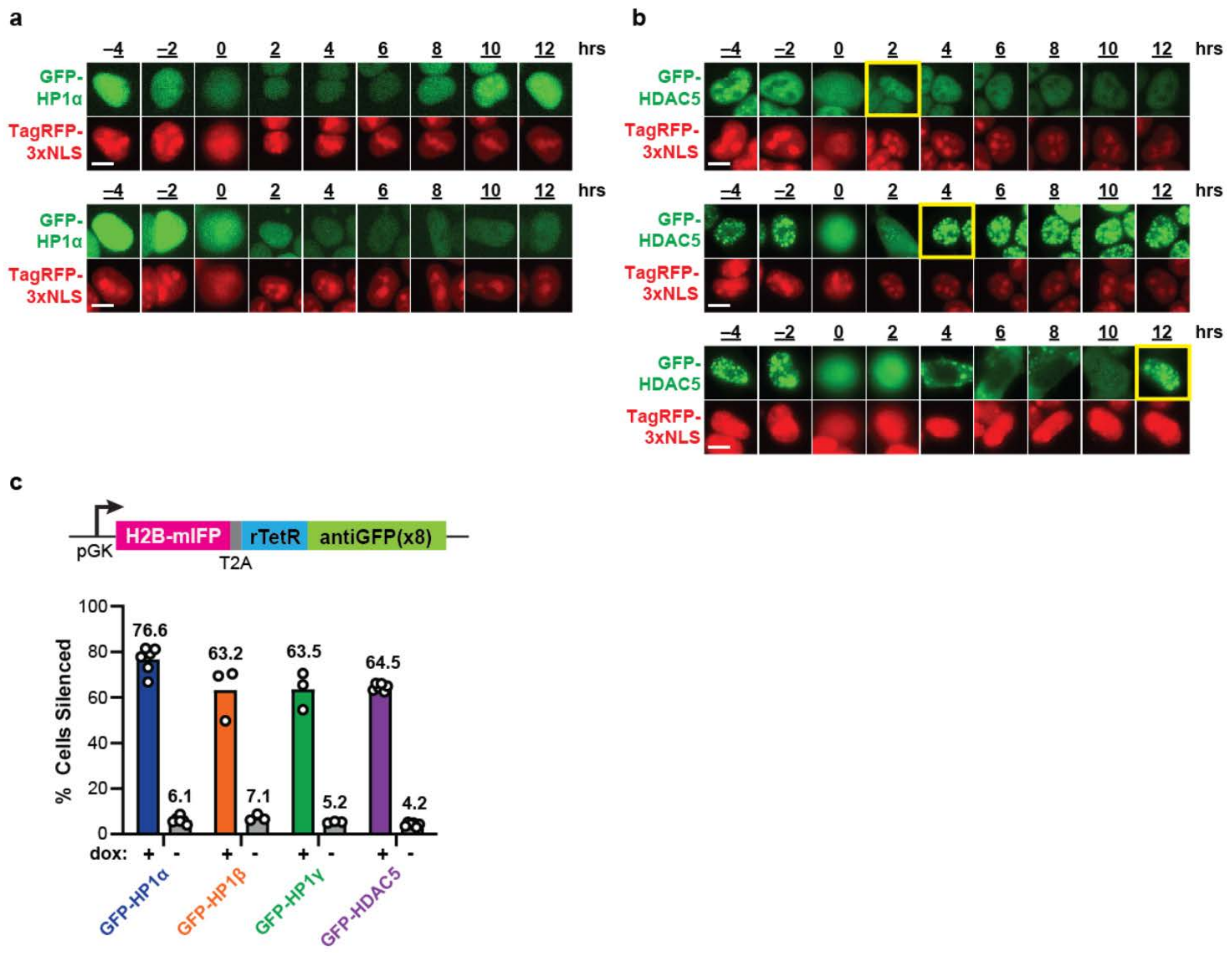


**Figure S1: Localization dynamics of GFP-tagged chromatin regulators and recruitment of multiple GFP-tagged chromatin regulators**

Time-lapse images of cells transiently expressed with **(a)** GFP-tagged HP1α and **(b)** HDAC5. Cells undergoing cell division are represented at time 0 hours. Yellow boxes highlight the re-entry of GFP-tagged chromatin regulators into the nucleus. White scale bars represent 10 μm. (**c**) Percent cells with reporter silenced after recruitment of an 8x repeat array of antiGFP nanobodies (+dox for 5 days). The experimental set up is the same as in Fig. 1a&d, with the reporter and nanobody array constructs stably integrated, and GFP-CRs transiently transfected. Each dot is an independently transfected biological replicate (GFP-HP1α: n = 6; GFP-HP1β: n = 3; GFP-HP1γ: n = 3; GFP-HDAC5: n = 6).

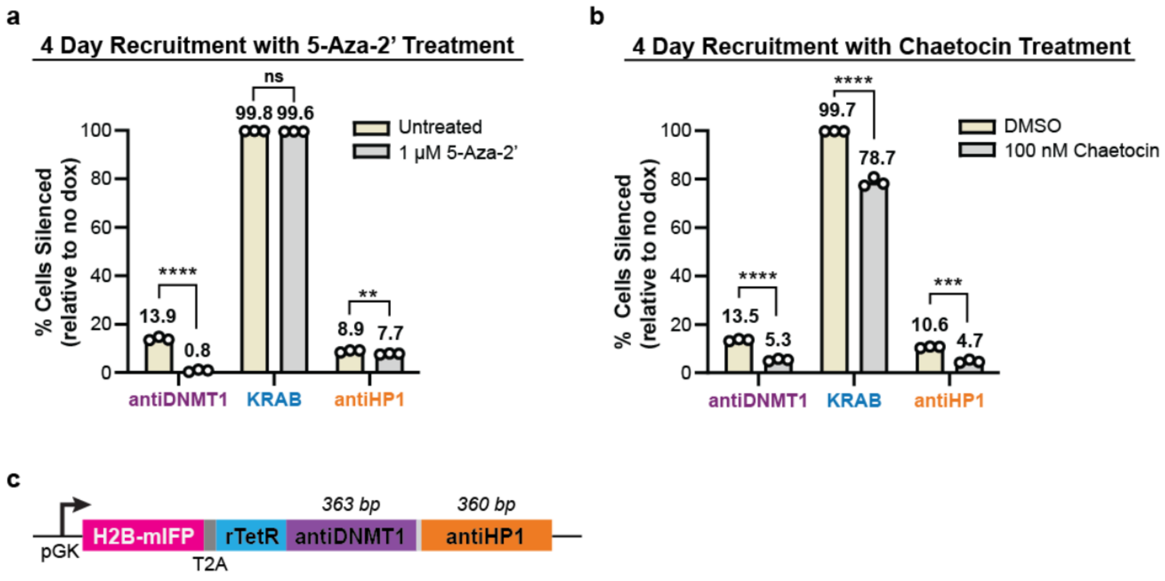


**Figure S2: Treatment of cells with DNA or histone methylation inhibitors**

Graphs comparing the percentage of cells with TagRFP silenced after four days of recruitment with doxycycline and **(a)** 5-aza-2'-deoxycytidine (5-Aza-2'), a DNA methyltransferase inhibitor, or **(b)** chaetocin, a broad-spectrum inhibitor of lysine histone methyltransferases. Means are from 3 replicates; statistical analysis by two-sample unpaired *t*-test. (**c**) Expression vector for H2B-mIFP and the rTetR-antiDNMT1-antiHP1 fusion under a pGK constitutive promoter with sizes of the DNA encoding for the nanobodies shown in base pairs (bp).

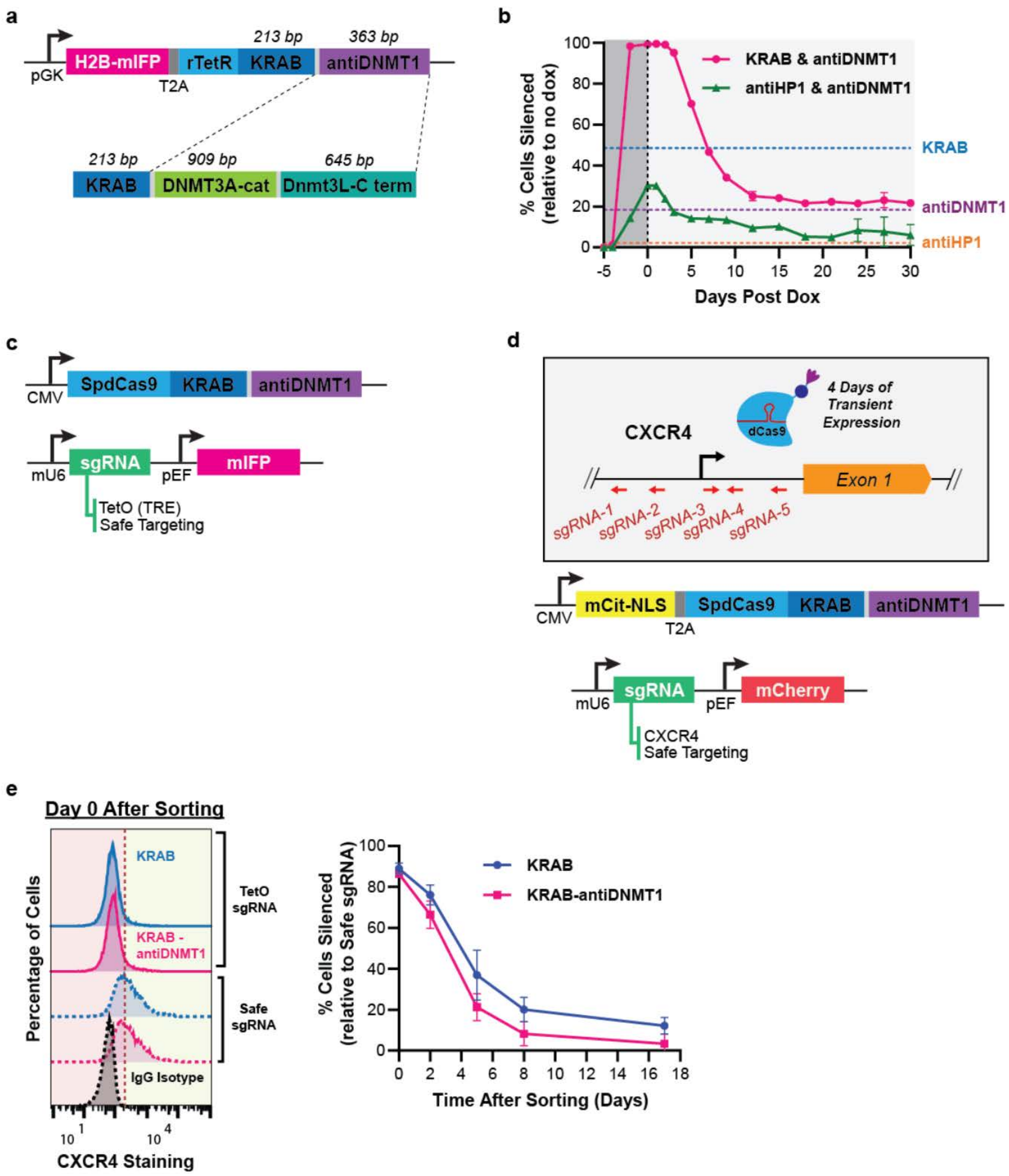


**Figure S3: Expression vectors and separate co-recruitment of regulators to the reporter gene for silencing**

(**a**) Expression vector for rTetR-KRAB-antiDNMT1 compared to the previously published KRAB-DNMT3A-3L fusion. (**b**) Percent cells with reporter silenced (relative to no dox controls) after co-recruitment of separate fusion of rTetR-effectors at the TagRFP reporter gene. Experimental setup is the same as in Fig. 2a&c. Included for reference are the percentages of cells permanently silenced after individual recruitment with KRAB, antiDNMT1, or antiHP1 (dashed lines) taken from Fig. 2d. (**c**) Expression vector for dCas9-KRAB-antiDNMT1 under a CMV constitutive promoter. The sgRNA (targeting either the TetO site or a safe genomic site with no annotated function) was expressed on a different vector driven by the mouse U6 promoter and contained a mIFP marker. (**d**) Diagram of sgRNA binding sites for the targeting of dCas9-KRAB-antiDNMT1 to the endogenous *CXCR4* gene. *CXCR4* is a cell surface transmembrane protein, which enables us to use conjugated fluorescent antibodies with flow cytometry to quantify gene expression. We cloned 5 sgRNAs (Table S1) spanning the upstream region of the transcriptional start site of this gene, targeting either the template or non-template strand. The dCas9 and sgRNA constructs were modified to express mCitrine and mCherry, respectively, to allow for cell sorting. (**e**) After transient expression and targeting at the endogenous *CXCR4* gene for 4 days, cells were sorted for the presence of both dCas9 (mCitrine positive) and sgRNA (mCherry positive). Cells were then immunostained for *CXCR4* expression and analyzed by flow cytometry (left). Means and standard deviations of percent cells with silent *CXCR4* from 2 replicates are shown throughout 17 days after sorting (right). Statistical analysis by two-sample unpaired *t*-test demonstrate no significant difference between the two effectors at all tested time points.

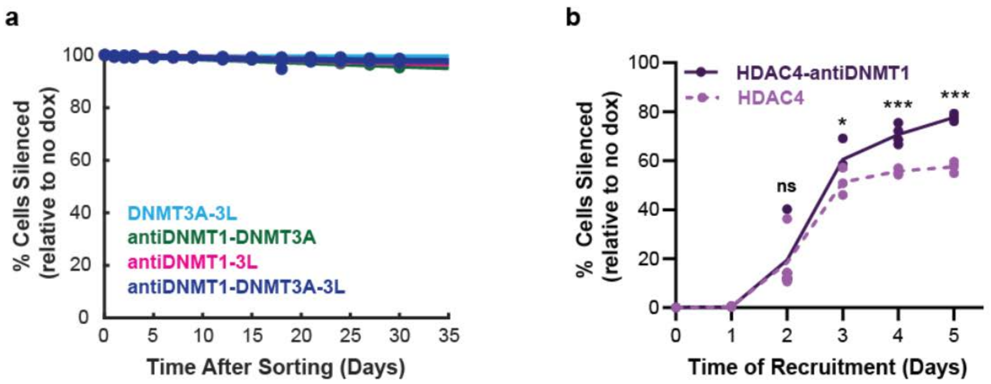


**Figure S4: Fusion of antiDNMT1 nanobody to DNMT3A-3L and HDAC4**

(**a**) After the rTetR-based DNA methyltransferases combinations were transiently transfected and treated with dox for 5 days, silenced cells were sorted, and reactivation was measured by flow cytometry throughout 30 days. Each dot is a biological replicate (DNMT3A-3L: n = 1; antiDNMT1-DNMT3A: n = 1; antiDNMT1-DNMT3L: n = 1; antiDNMT1-DNMT3A-3L: n = 3) and the data are fitted with an exponential decay curve. (**b**) Transient expression and recruitment of the rTetR-HDAC4-antiDNMT1 fusion to the reporter gene throughout 5 days (n = 4 replicates; statistical analysis by two-sample unpaired *t*-test).

**Table S1: List of sgRNA Sequences**

Sequences of sgRNAs used in this study are listed below. For endogenous gene targeting of *CXCR4*, sgRNA naming is based on the binding coordinates relative to the transcription start site (TSS), and whether they target the template (T) or non-template (NT) strand. A negative coordinate indicates a binding location upstream of the TSS and a positive coordinate indicates a binding location downstream of the TSS.

| **Target Gene** | **sgRNA Sequence** | **Notes** |
|---|---|---|
| TetO_sgRNA | GTACGTTCTCTATCACTGATA | Targets the 5x tetracycline operator (TetO) sequence. |
| Safe_sgRNA-1 | GATAGGCACAGGAAATTTGG | Safe guide control used when targeting the reporter gene. |
| Safe_sgRNA-2 | GCACATTTGGATTTCATGTC | Safe guide control used when targeting the *CXCR4* gene. |
| *CXCR4*_sgRNA-1 (NT, -204) | GAGGCATTTCCTAAGTTTGA | |
| *CXCR4*_sgRNA-2 (NT, -126) | GCGCGGCTTGGGAAGCCCAG | |
| *CXCR4*_sgRNA-3 (T, +36) | CAGGTAGCAAAGTGACGCCGA | Previously validated in Yeo et al. 2018. |
| *CXCR4*_sgRNA-4 (NT, +102) | AACCGCTGGTTCTCCAGATG | |
| *CXCR4*_sgRNA-5 (NT, +501) | TTCCCCAAGGAAGAGACCGG | |

**Supplementary Text: Modeling of Gene Control**

Gene silencing and memory upon recruitment of antiDNMT1 at a gene can be described by a kinetic model consisting of 3 gene states (Fig. 4d). The time evolution for the fraction of cells in each of the three states - active (A), reversibly silent (R), and irreversibly silent (I) - can be calculated from the set of differential equations associated with the kinetic rates in Fig. 4d, during recruitment and release, respectively. We denoted the cells in each state with a "+" subscript (i.e. $A_+$), to indicate when the equations describe the "+dox" silencing period, and with a "-" subscript for the "-dox" period (i.e. $A_-$). At the end, we combine them into a single function that describes the behavior across the two periods (no subscript, i.e. $A$).

**Derivation of the fractions of cells in each gene state during antiDNMT1 recruitment (+dox)**

During recruitment, we denote the time of dox induction as *t*, and can describe the fraction of cells in each state according to the kinetic model in Fig. 4d, top:

$$\frac{dA_+}{dt} = -k_S A_+ \qquad \frac{dR_+}{dt} = k_S A_+ - k_I R_+ \qquad \frac{dI_+}{dt} = k_I R_+$$

We can solve for the fraction of cells active during recruitment directly from the first equation above:

$$A_+(t) = A_0 e^{-k_S t},$$

where $A_0$ is the fraction of cells active at the beginning of recruitment. Since the total fractions of silent and active cells must add up to 1, the total fraction of silent cells (S) can be described as: $S_+(t) = 1 - A_0 e^{-k_S t}$. This equation is sufficient to describe and fit the silencing phase. Note that in order to express percentages of cell silent as a function of time (as presented in the main figures), we can simply multiply each equation by 100.

To understand how many cells are committed to long term memory (and thus the behavior after CR release), we need to know how silent cells partition into reversibly versus irreversibly silent over time: To solve for $R_+(t)$, we replace $A_+(t) = A_0 e^{-k_S t}$, into $dR_+/dt = k_S A_+ - k_I R_+$ :

$$\frac{dR_+}{dt} = k_S A_0 e^{-k_S t} - k_I R_+.$$

We can solve this differential equation to get:

$$R_+(t) = \frac{k_S}{k_I - k_S} A_0 (e^{-k_S t} - e^{-k_I t}) + R_0 e^{-k_I t}$$

The fraction of irreversible silenced cells over time is: $I_+(t) = 1 - A_+(t) - R_+(t)$.

In summary, the function describing the fraction of cells in each of the three states for a given silencing dox signal of duration $t$ (and no reactivation period):

$$A_+(t) = A_0 e^{-k_S t}$$

$$R_+(t) = \frac{k_S}{k_I - k_S} A_0 (e^{-k_S t} - e^{-k_I t}) + R_0\, e^{-k_I t} \qquad (1)$$

$$\mathrm{I}_+(t) = 1 - A_+(t) - \mathrm{R}_+(t) = 1 - \frac{k_I}{k_I - k_S} A_0 e^{-k_S t} + (\frac{k_S}{k_I - k_S} A_0 - R_0)\, e^{-k_I t}$$

**Derivation of the fractions of cells in each gene state during antiDNMT1 release (-dox)**

During release (-dox), we denote the time since dox was removed as $\tau$, and can describe the fraction of cells in each state according to the kinetic model in Fig. 4d, bottom:

$$\frac{dA_-}{d\tau} = k_A R_- \qquad \frac{dR_-}{d\tau} = -k_A R_- \qquad \frac{dI_-}{d\tau} = 0$$

We can solve for the fraction of cells reversibly silent during release from the $dR_-/dt$ equation above:

$$R_-(\tau) = R_1 e^{-k_A \tau},$$

where $R_1$is the fraction of reversibly silenced cells at the beginning of release (end of recruitment).

During release, the irreversible fraction stays constant, at the value reached at the end of recruitment, $I_1$: $\quad I_-(\tau) = I_1$

The total fraction of cells silent over time in the release phase, $S_-(\tau) = R_-(\tau) + I_-(\tau)$, is described by an exponential decay to the irreversible fraction:

$$S_-(\tau) = R_1 e^{-k_A \tau} + I_1.$$

The fraction of cells active is: $A_-(\tau) = 1 - I_-(\tau) - R_-(\tau)$.

In summary:

$$\begin{aligned} A_-(\tau) &= 1 - I_1 - R_1 e^{-k_A \tau} \\ R_-(\tau) &= R_1 e^{-k_A \tau} \\ I_-(\tau) &= I_1 \end{aligned} \qquad (2)$$

**General solution and additional considerations for fitting the data**

We can combine the results of the two previous sections to obtain general solutions that describe the cells in each state across both recruitment and release times. The starting fractions of cells $R_1$, $I_1$, at the beginning of the –dox release, depend on the duration of +dox period $t$: $R_1 = R_+(t)$, and $I_1 = I_+(t)$. Therefore, based on the solutions derived in the release section above (equations (2)):

$$A(t,\tau) = 1 - I_+(t) - R_+(t) e^{-k_A \tau}$$

$$R(t,\tau) = R_+(t)\, e^{-k_A \tau}$$

$$I(\mathrm{t},\tau) = \mathrm{I}_+(\mathrm{t})$$

By replacing $A_+(t)$ , $R_+(t)$ , and $I_+(t)$ with the set of equations (1) derived in the recruitment section, the general solution after a recruitment signal (+dox) of duration *t*, and reactivation period (-dox) of duration τ becomes:

$$A(t,\tau) = \frac{k_I}{k_I - k_S} A_0 e^{-k_S t} - (\frac{k_S}{k_I - k_S} A_0 - R_0)\, e^{-k_I t} - \left(\frac{k_S}{k_I - k_S} A_0 (e^{-k_S t} - e^{-k_I t}) + R_0\, e^{-k_I t}\right) e^{-k_A \tau}$$

$$R(t,\tau) = \left(\frac{k_S}{k_I - k_S} A_0 (e^{-k_S t} - e^{-k_I t}) + R_0\, e^{-k_I t}\right) e^{-k_A \tau} \qquad (3)$$

$$I(\mathrm{t},\tau) = 1 - \frac{k_I}{k_I - k_S} A_0 e^{-k_S t} + (\frac{k_S}{k_I - k_S} A_0 - R_0)\, e^{-k_I t}$$

During recruitment, we observe a time lag between dox addition and the onset of silencing ($T_{lag}$). Therefore, for fitting purposes, in the equations above $t$ becomes $t - T_{lag}$, for recruitment times larger than $T_{lag}$, and no changes in the fractions of silent/active cells are allowed at shorter times.

The final equations used to calculate the fraction of cells silent over time during a period of +dox induction are:

$$A_+(t) = \begin{cases} A_0 & \text{, for } t < T_{lag} \\ A_0 e^{-k_S(t - T_{lag})} & \text{, for } t \geq T_{lag} \end{cases}$$

$$R_+(t) = \begin{cases} R_0 & \text{, for } t < T_{lag} \\ \frac{k_S}{k_I - k_S} A_0 (e^{-k_S(t-T_{lag})} - e^{-k_I(t-T_{lag})}) + R_0\, e^{-k_I(t-T_{lag})} & \text{, for } t \geq T_{lag} \end{cases}$$

$$\mathrm{I}_+(\mathrm{t}) = \begin{cases} 1 - A_0 - R_0 & \text{, for } t < T_{lag} \\ 1 - \frac{k_I}{k_I - k_S} A_0 e^{-k_S(t-T_{lag})} + (\frac{k_S}{k_I - k_S} A_0 - R_0)\, e^{-k_I(t-T_{lag})} & \text{, for } t \geq T_{lag} \end{cases}$$

For fitting a continuous dox signal and the reactivation following it, we start with all cells silenced: $A_0 = 1, R_0 = 0, I_0 = 0$. Therefore, the equations for silencing simplify to:

$$A_+(t) = \begin{cases} 1 & \text{, for } t < T_{lag} \\ e^{-k_S(t - T_{lag})} & \text{, for } t \geq T_{lag} \end{cases}$$

$$R_+(t) = \begin{cases} 0 & \text{, for } t < T_{lag} \\ \frac{k_S}{k_I - k_S} (e^{-k_S(t-T_{lag})} - e^{-k_I(t-T_{lag})}) & \text{, for } t \geq T_{lag} \end{cases}$$

$$\mathrm{I}_+(\mathrm{t}) = \begin{cases} 0 & \text{, for } t < T_{lag} \\ 1 - \frac{k_I}{k_I - k_S} e^{-k_S(t-T_{lag})} + \frac{k_S}{k_I - k_S}\, e^{-k_I(t-T_{lag})} & \text{, for } t \geq T_{lag} \end{cases}$$

The general solution in the case when we start with all cells active ($A_0 = 1, R_0 = 0, I_0 = 0$) becomes:

$$A(t,\tau) = \frac{k_I}{k_I - k_S} e^{-k_S(t-T_{lag})} - \frac{k_S}{k_I - k_S}\, e^{-k_I(t-T_{lag})} - \frac{k_S}{k_I - k_S} (e^{-k_S(t-T_{lag})} - e^{-k_I(t-T_{lag})}) e^{-k_A \tau},$$

$$R(t,\tau) = \frac{k_S}{k_I - k_S} (e^{-k_S(t-T_{lag})} - e^{-k_I(t-T_{lag})})\, e^{-k_A \tau}\,, \qquad (4)$$

$$I(t,\tau) = 1 - \frac{k_I}{k_I - k_S} e^{-k_S(t-T_{lag})} + \frac{k_S}{k_I - k_S}\, e^{-k_I(t-T_{lag})}\,,$$

for all recruitment times $t \geq T_{lag}$, and $A(t,\tau) = 1, R(t,\tau) = 0\,, I(t,\tau) = 0$ for $t < T_{lag}$.
